# Continuous mass photometry by single molecule trapping

**DOI:** 10.64898/2026.02.02.703204

**Authors:** Dan Loewenthal, Roi Asor, Stephen Thorpe, Jack S. Peters, Konstantin Zouboulis, Merve Kaplan, Jan Christoph Thiele, Jack Bardzil, Seham Helmi, Sean A. Burnap, Tiong Kit Tan, Weston B. Struwe, Justin L.P. Benesch, Philipp Kukura

## Abstract

The dynamic choreography of biomolecular interactions underpins the processes of life, but its direct observation remains challenging. Here, we introduce confined diffusion mass photometry, enabling hour-long, mass-resolved observation of individual biomolecules, their complexes and interactions with up to sub-kDa mass precision and ms temporal resolution. Our approach represents a quantitative time-resolved single-molecule measurement modality for studying complex biomolecular mechanisms in action.

## Main

Technologies capable of studying single biomolecules have transformed our understanding of biomolecular processes^1-3^. Nevertheless, one of the core promises of single molecule biophysics – quantitative, time-resolved observation of biomolecular interactions at the single-molecule level - has remained challenging. Achieving this would reveal information key to biological understanding and therapeutics development, yet difficult to access in ensemble measurements: the sequence of biomolecular interactions, dwell time distributions, and transitions that define biomolecular mechanisms.

Single-molecule mass measurement by light scattering^4-7^ promises a direct and quantitative way to study biomolecular interactions in a time-resolved manner. This is because mass is a universal and specific observable, directly reporting on biomolecular composition, while light scattering, being unaffected by photobleaching, is effectively time-unlimited, thus well-suited for extended, time-resolved studies. Despite this promise, current approaches are constrained in observation time, operation in biologically-relevant concentrations, or both. Cavity-enhanced methods^5^ achieve remarkable sensitivity, but are limited to the microsecond regime determined by cavity transition times. Nanofluidic approaches^6^ extend observation times in free solution, but demand very low analyte concentrations and have yet to demonstrate interactions or resolution of heterogeneous mixtures. Mass photometry relies on destructive binding to a glass surface^8-10^, with observation limited to a few seconds at best as biomolecules diffuse on a supported lipid bilayer (SLB)^11,12^.

We reasoned that combining single-molecule trapping using confined SLBs with mass photometry could overcome these limitations. In this approach, individual molecules remain in the microscope’s field of view, avoid co-localisation with other trapped biomolecules, and can freely interact with other biomolecules from solution while their mass is continuously monitored.

We start by photolithographically preparing an SLBs geometrically confined by dense poly-L-lysine polyethylene glycol (PLL-PEG) brushes (**Fig. 1a,b**), with a low density of capture-functionalised lipids (e.g. NTA)^13-17^. Here, both the SLB and PEG provide excellent surface passivation except for the species of interest, allowing the presence of other biomolecules in solution. Following the addition of the target protein, in this case 6xHis-tagged SARS-CoV-2 spike glycoprotein trimers, we find anywhere between zero and three spike glycoproteins bound to SLB traps (**Fig. 1c, SFig. 1**)

**Figure 1.**
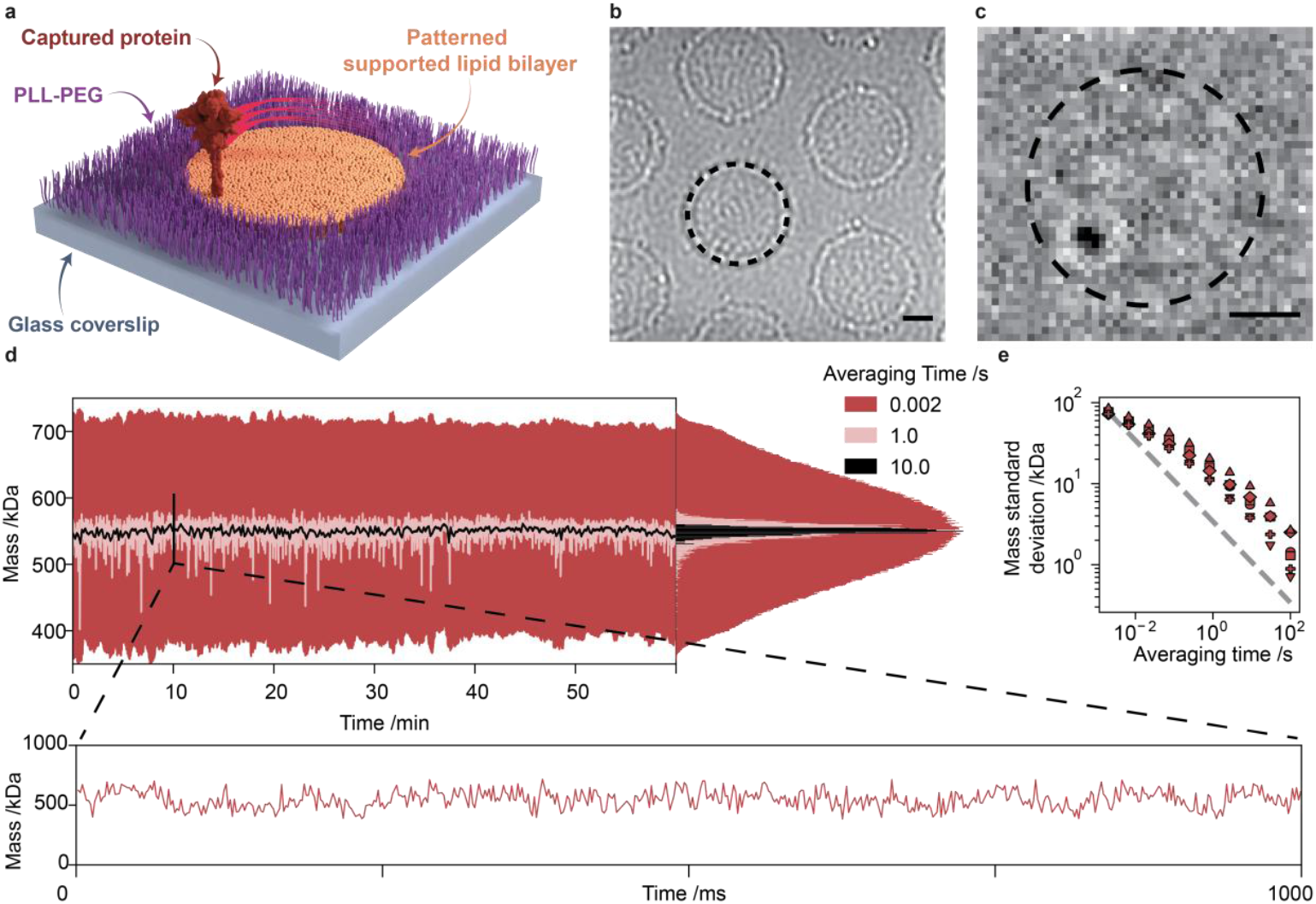
Hour long mass monitoring of a single protein. **(a)** Schematic of a Confined Diffusion Mass Photometry (CDMP) experiment. **(b)** Mass photometry image of confined supported lipid bilayer traps. **(c)** Median processed mass photometry image revealing a diffusing protein. Scale bars: 1 µm. **(d)** Top: CDMP mass vs time trace of a single diffusing SARS-CoV-2 spike glycoprotein. Bottom: One second zoom at 0.002 s averaging time. **(e)** Mass precision as a function of averaging time for 6 different proteins.

Using commercial MP instruments (wavelength <530 nm), we observed that the diffusing species became immobilised (**SFig. 2**), likely due to defect formation in the patterned SLB, terminating our measurement. Upon moving to 638 nm illumination, however, we could frequently monitor individual proteins for up to an hour before immobilisation (**SFig. 3-5**). For an imaging frame rate of 497 Hz, an hour-long trajectory thus consists of ∼1.8 million position and mass measurements, from which we can extract both mobility and molecular mass **(Fig. 1d, SFig. 3-5**, and **Supplementary Video 1**). Moreover, as a direct consequence of extended observation, we can improve mass precision by signal averaging at the expense of temporal resolution,^18,19^ reaching ∼1 kDa mass precision with 100 s temporal averaging (**Fig. 1e**).

To assess the quantitative ability of CDMP in detecting mass changes caused by biomolecular binding, we measured the interactions between SARS-CoV-2 spike trimers and a series of ligands with progressively smaller masses: IgG antibodies (150 ± 1 kDa), IgG Fab fragments (49 ± 1 kDa), and nanobodies (16,480 ± 1 Da). In standard MP (**Fig. 2a**), distinct molecular species can be resolved for the antibody interaction. However, for the Fab and nanobody (**Fig. 2b**), these species become unresolvable, as the measurement error for the masses of the coexisting species is comparable in magnitude to the mass spacing between them.

**Figure 2.**
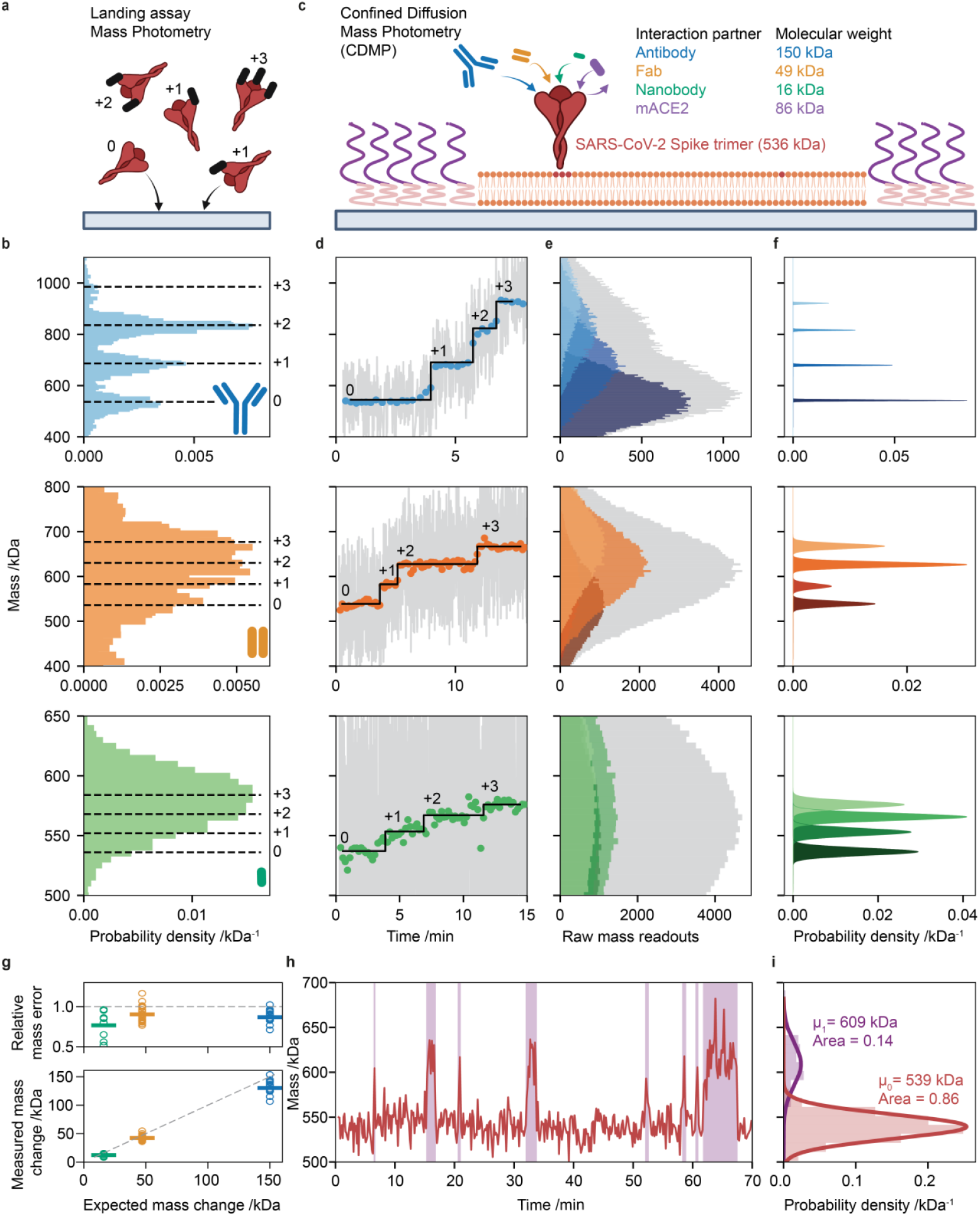
Real-time monitoring of single-molecule biochemical reactions through step-changes in mass. **(a)** Schematic of a standard MP landing assay. **(b)** MP histograms for SARS-CoV-2 spike mixtures (Top: antibody, middle: fab, bottom: nanobody). **(c)** Schematic of the CDMP experiment. **(d)** CDMP mass vs time traces (grey: raw, scatter: 10 second averaged) of a single confined SARS-CoV-2 spike protein binding to: Top: antibody. Middle: fab. Bottom: nanobody from solution. **(e)** Raw (grey) and state-specific (coloured) CDMP raw mass histograms. **(f)** Corresponding CDMP probability density histograms. **(g)** Measured vs expected mass changes and relative mass error for CDMP. **(h)** CDMP mass vs time trace of a SARS-CoV-2 spike protein interacting transiently with a monomeric ACE2 protein. **(i)** Gaussian fits to the states found in (h).

For trapped molecules (**Fig. 2c**), however, a single molecule occupies different molecular states over time, and these states appear sequentially rather than simultaneously. Thus, the challenge now becomes one of detecting changes in molecular mass, as opposed to resolving a mixture **(Fig. 2d)**. By tuning the averaging time, we can reliably distinguish successive states.

This principle is analogous to super-resolution fluorescence microscopy, where fluorophores are switched on and off at different times so they can be localized independently. Once states are identified, their average mass and variability can be used to construct mass-probability spectra (**Fig. 2e,f**). In each case, we observe a maximum of three binding events, corresponding to the three binding sites of the spike trimer (**Fig. 2d-f, SFig. 6-8)**. The associated mass changes scale well with the expected mass of the ligand, subject to a ∼15% offset consistent with an expected reduction in interferometric contrast stemming from the distance of the receptor binding domain of the spike trimer from the SLB^20^ (**Fig. 2g, SFig. 9**). The chosen interaction partners exhibit tight binding associated with a slow off-rate, directly visible from the mass vs time traces showing no evidence for ligand unbinding on the >10 min timescale.^21,22^ To demonstrate dynamic (un)binding, we chose the interaction with a monomeric variant of the spike receptor, ACE2 (angiotensin converting enzyme 2) (mACE2). mACE2 was chosen to challenge the technique, featuring lower mass and faster unbinding kinetics compared to dimeric ACE2. Individual traces now show binding followed by unbinding events (**Fig. 2h,i, SFig. 10**), in good agreement with the expected mass change of mACE2 with a stochastic range of dwell times.

Currently, our method can reliably track dynamics where the analyte starting mass is >200 kDa, with mass changes of >20 kDa and lifetimes on the order of seconds to tens of minutes. We anticipate rapid developments in all key aspects of this method, further expanding the scope of the technique. Larger measurement fields of view and optimised trap geometries will improve throughput by allowing more than one protein to be measured simultaneously. Optimisation of supported lipid bilayer chemistry and laser pulsing will lower background and measurement noise, cutting the required averaging time for kDa precision and enabling even longer observation times. Most excitingly, the use of a capture complex on the order of 200 kDa bridging the lipid and protein will effectively eliminate the detection limit of mass photometry, now only bound by mass precision rather than background noise. Taken together, we show that our platform enables quantitative, real-time observation of biomolecular interactions at the single-molecule level.

## Acknowledgements

D.L. is supported by a Clarendon scholarship, a Menasseh Ben Israel scholarship, and a Kingsgate scholarship. R.A. was supported by EMBO long-term postdoctoral fellowship ALTF-198-2020. S.T. is supported by the Biotechnology and Biological Sciences Research Council BB/W00349X/1. J.S.P. was supported by the ERC grant Photomass (819593). K.Z. was supported by the Wellcome Trust LEAP programme NanoQuest. M.K. is supported by the Study Abroad Program of the Ministry of National Education, The Republic of Türkiye. J.C.T. is supported by a Schmidt Sciences LLC. J.B. is supported by an EPRSC Doctoral Training Partnership. S.H. is supported by the EPSRC (EP/T03419X/1) and a BBSRC Transformable Research Technologies (UKRI1877). S.A.B. is supported by a Wellcome Early-Career Award 312750/Z/24/Z. T.K.T is supported by a Coalition of Epidemics Preparedness Innovation grant under the broadly protective coronavirus vaccine (BPCV) portfolio. W.S. is supported by funding a UK Research and Innovation Future Leaders Fellowship (MR/V02213X/1). J.L.P.B. is supported by the BBSRC (BB/W00349X/1). P.K. is supported by the EPSRC (EP/T03419X/1 and EP/W001055/1)

## Author contributions

P.K., D.L., and R.A. conceptualized and designed the research; D.L. performed the experiments; R.A. and D.L. wrote the code and analysed the data; D.L. and S.T. developed the photolithography procedure; J.S.P., J.C.T., J.B., D.L., and P.K. designed and built the custom optical setup; S.H. assisted with initial experiments; K.Z., M.K., and S.A.B. supplied ACE2, spike, and nanobody samples; T.K.T. supplied spike antibody and Fab samples, W.B.S., J.L.P.B. and P.K. supervised the research; D.L., R.A. and P.K. wrote the paper.

## Methods

### Stocks and reagents

Isopropanol (24137-M), silicone gaskets (GBL 103280), HEPES (H3375), magnesium chloride hexahydrate (M2670) and KCl (P9541) were purchased from Merck Life Science UK Limited. SARS-CoV-2 nanobodies were bought from Nanotag biotechnologies (Cat No N3505). Glass slides (11836933) were purchased from Fisher Scientific. Lipids (1-palmitoyl-2-oleoyl-sn-glycero-3-phosphocholine, POPC: 850457, 1,2-dioleoyl-sn-glycero-3-phosphoethanolamine-N-[methoxy(polyethylene glycol)-550], DOPE-PEG550: 880530C, 1,2-dioleoyl-sn-glycero-3-[(N-(5-amino-1-carboxypentyl)iminodiacetic acid)succinyl] (nickel salt), DGS-NTA(Ni): 790404P) were purchased from Avanti Polar Lipids. PLL(20)-g[3.5]-PEG(2) was purchased from SuSoS surface technology.

### Liposome preparation

Similarly to our previously described protocol^11^, liposomes were prepared by dissolving POPC, DOPE-PEG550, and DGS-NTA(Ni) in chloroform and mixing them in a clean glass tube at a 97–2-1 molar ratio (500 μM total lipid concentration). The mixture was then dried under a constant nitrogen stream via rotary evaporation and further dried under vacuum for 1 h at room temperature (22–23 °C). A total of 500 μl HKS-150 (20 mM HEPES, pH 7.4, 150 mM KCl) was added to the dried lipids and the mixture was covered with parafilm, incubated at 50 °C in a water bath for 1 h, briefly vortexed, and stored overnight at room temperature. The resulting liposome mixture was transferred into a 1.5 ml Eppendorf tube and kept in ice-water and sonicated using a 3 mm probe at 25% amplitude with a 1 s on–3 s off sonication cycle (Sonics & Materials) for 10 min (that is, 40 min in total). Sonicated liposomes were then spun at 21,130 ×g for 30 min at 4 °C and the supernatant was collected in an Eppendorf tube, stored at 4 °C, and used within 30 days.

### Patterned supported lipid bilayers

Confined supported lipid bilayers were made based on a published protocol with some adjustments^17^. Briefly, glass coverslips were placed in a metal rack and cleaned by sonication for 5 min in Milli-Q water followed by sonication for 5 min in a solution of 50/50 Milli-Q water/isopropanol. The coverslips were then dried with nitrogen then plasma cleaned using a Zepto plasma cleaner (Diener electronic) operating at 40% power for 5 minutes. The coverslips were then rinsed with Milli-Q water and dried with nitrogen. The coverslips were then fitted with gaskets and were incubated with a 2 μg/mL PLL(20)-g[3.5]-PEG(2) solution in Milli-Q water for 30 min. Before being subjected to photolithography, coverslips were thoroughly rinsed with Milli-Q water and dried with nitrogen. The coverslips were then exposed to deep UV light in a mask aligner (Suss MJB4, HgXE 500W source) for 60 min through a home-made photolithography mask. The coverslips were then thoroughly rinsed with Milli-Q and dried with nitrogen. These were kept in -20 °C for up to three months before use. Before every experiment the coverslips were brought into room temperature then exposed to 25 μL fusion buffer (20 mM HEPES, pH 7.4, 150 mM KCl, 2 mM MgCl) and 5 μL of liposome solution in a silicone gasket. After bilayer formation (monitored by mass photometry), which took around 5-10 minutes the gaskets were washed with PBS five times by replacing the full volume of solvent in the gasket. The surfaces were blocked with 0.1 mg/mL PLL-PEG for 30 minutes unless otherwise noted and then washed again ten times with PBS by solvent volume replacement.

### Confined diffusion mass photometry optical setup

Measurements were performed on a home-built mass photometry setup equipped with a red diode laser (Lasertack, 638 nm, 1W) and a Blackfly 17S7M-C, FLIR camera. Where acoustooptic deflector (AOD) scanning was controlled using a Digilent Analog Discovery 2 device (Digilent). The setup was operated using a home-built LabView script.

### Data acquisition

#### Confined diffusion mass photometry

Mass photometry images were measured in a field of view (FOV) of approximately 6×6 μm which was centred on a ∼3.5 μm diameter trap. The area was continuously scanned and images were taken using an exposure time of 1700 μs at 497 Hz for the long tracking experiments. For the step change and transient binding experiments the AOD scanning was pulsed to position the beam further from the trap when the camera is not exposed and the frame rates were 250 Hz and 10 Hz respectively with the same exposure time per frame.

#### CDMP - Long tracking experiments

Supported lipid bilayers were not blocked before protein addition. Spike was added at a 5 nM concentration in solution for capture and washed within 30 seconds for a capturing occupancy of zero to two particles per confined SLB. The trap was illuminated in the CDMP setup continuous illumination at a 497 Hz frame rate and 1700 μs exposure time.

#### CDMP - Step change experiments

Spike trimers were added at a 5 nM concentration in solution for capture and washed within 30 seconds for a capturing occupancy of zero to two particles per confined SLB. The trap was illuminated in the CDMP setup at pulsed illumination at a 250 Hz frame rate and 1700 μs exposure time. For the first three minutes in each experiment no interaction partner was added in order to establish a baseline mass. At the three-minute mark an antibody \ fab \ nanobody was added. Antibodies were added as 100 nM by a 0.75:99 dilution from a 13.3 μM stock, every five minutes the solution was mixed with a pipette. Fabs were added as 100 nM by a 10:90 dilution from a 1 μM stock then every five a further 10 μL of the fab stock were added, increasing the concentration to facilitate binding within the measurement period. Nanobodies were added as 100 nM by a 10:90 dilution from a 1 μM stock then every five a further 10 μL of the nanobody stock were added, increasing the concentration to facilitate binding within the measurement period.

#### CDMP - Transient binding experiments

Spike trimers were added at a 5 nM concentration in solution for capture, and washed within 30 seconds for a capturing occupancy of zero to two particles per confined SLB. The trap was illuminated in the CDMP setup at pulsed illumination at a 10 Hz frame rate and 1700 μs exposure time. For the first three minutes in each experiment no interaction partner was added in order to establish a baseline mass. At the three-minute mark 210 nM of mACE2 were added while spike was continuously measured.

#### Landing assay mass photometry

Landing assay mass photometry measurements were performed on a Refeyn TwoMP at the “Regular” FOV (10.9 x 4.3 µm and 498 Hz). They were performed on microscopy slides that were cleaned by consecutive sonication in Milli-Q water followed by 50/50 Milli-Q water/isopropanol. The slides were fitted with silicone gaskets before measurement and the total volume of protein solutions was 20 μL. Interaction mixtures were incubated at equimolar concentrations at 25 nM for 10 minutes then measured as a 1:1 dilution with PBS in the mass photometry gasket for 60 seconds.

### Data processing

#### Confined diffusion mass photometry

Confined diffusion mass photometry measurements were analysed using an in-house python script^11,20^ adjusted for single trap analysis (**SFig. 11**). Briefly, proteins were detected using a Laplacian of gaussian threshold filter on a median processed mass photometry movie, this reveals the much smaller protein signal (∼0.3%, **Fig. 1c**) compared to the raw image, but demands the analyte to be mobile^11,12^. The detected proteins were then fitted an analytical PSF function^11^ calibrated to the home-built mass photometry setup to determine the protein contrast. The fitted events were then filtered for single protein analysis. Lastly, the starting mass was set as the mean landing assay mass (**SFig. 12,13**). The results were then presented as a running median of the raw fitted event masses or averaged mass values of consecutive time windows. For step detection, we implemented a step detection algorithm^23^ applying a maximum threshold of 10 steps per trajectory and minimum step size thresholds of 100,30 and 8 for the Antibody, fab and nanobody respectively. The transient binding experiments (mACE2) were time averaged to 0.2 Hz then fitted a two-state Hidden Markov Model.

#### Landing assay mass photometry

Landing assay mass photometry data was analysed using the DiscoverMP software (V2024 R1). The frames were further averaged to navg = 15 (33.2 Hz) using the advanced software settings. Results are represented as the mean value of three technical replicates.

## Data availability

The raw data required to reproduce all of the manuscript figures will be deposited in the University of Oxford Research Archive upon manuscript acceptance.

## Code availability

The python software and Jupyter notebooks required to reproduce all of the manuscript figures from the raw data are available upon request.

