## Supplementary Information for "Continuous mass photometry by single molecule trapping"

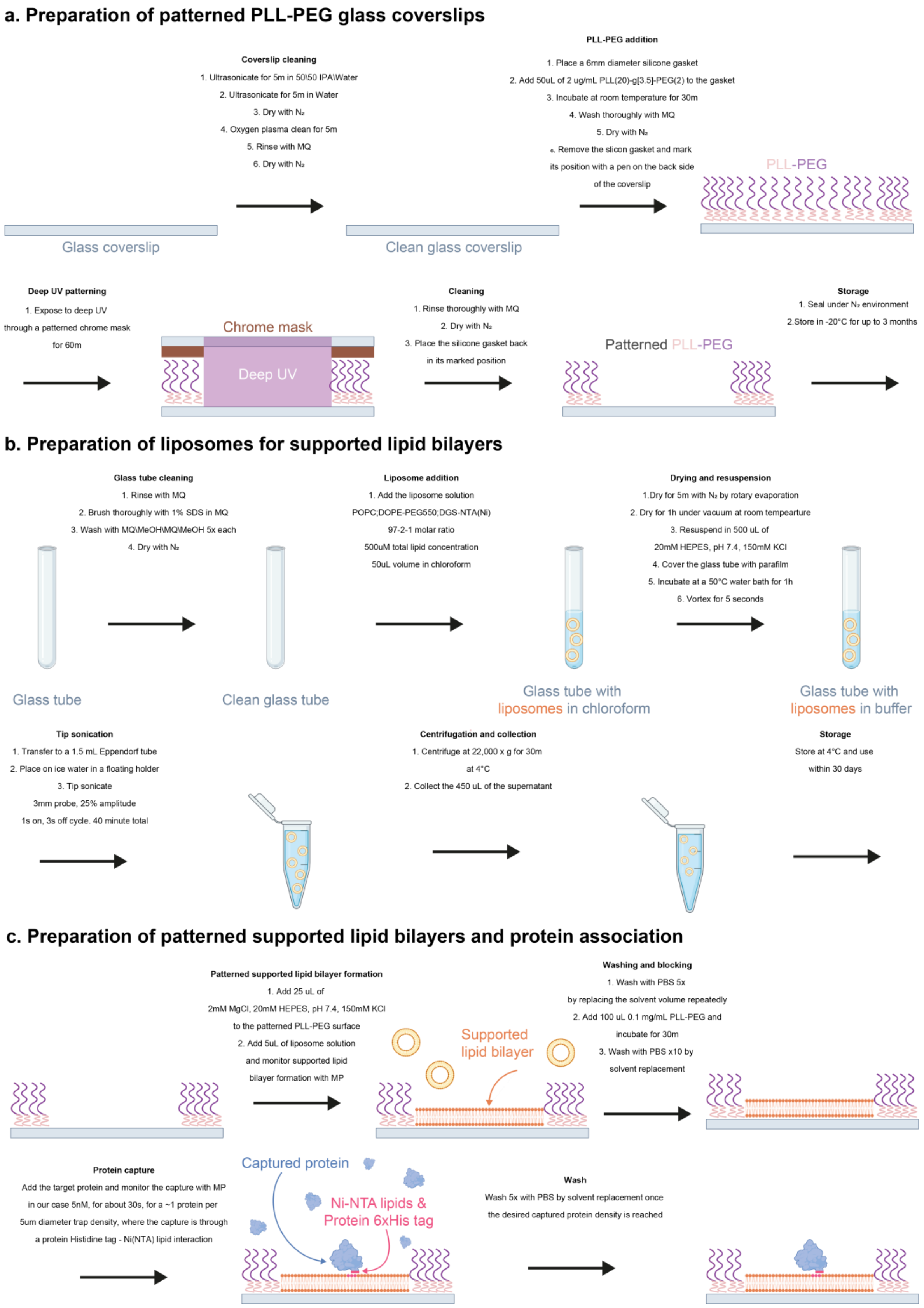

**Supplementary figure 1. Schematics for the experimental preparation of confined diffusion mass photometry experiments.**
**(a)** preparation of coverslips covered with patterned poly-L-lysine (PLL) – polyethylene glycol (PEG) through deep UV photolithography. **(b)** Preparation of liposomes solution for supported lipid bilayers. **(c)** preparation of patterned supported lipid bilayers and protein association.

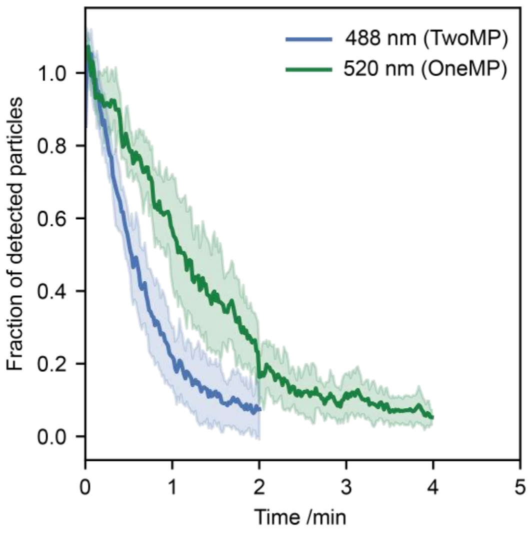

**Supplementary figure 2. Illumination dependent immobilisation of confined SLB captured SARS-CoV-2 spike glycoproteins**.

Spike glycoproteins were captured to a 10 x 10 μm$^2$ confined SLB and the number of detected (diffusing) proteins was monitored over time. Solid and transparent lines represent the mean value and standard deviation of technical replicates respectively (n=10 TwoMP, n=14 OneMP). Each replicate trace was normalised to the average number of particles detected in the first 10 seconds of measurement. The measured field of view sizes were 16.9x12.0 μm$^2$ for the TwoMP and 13.7x13.7 μm$^2$ for the OneMP.

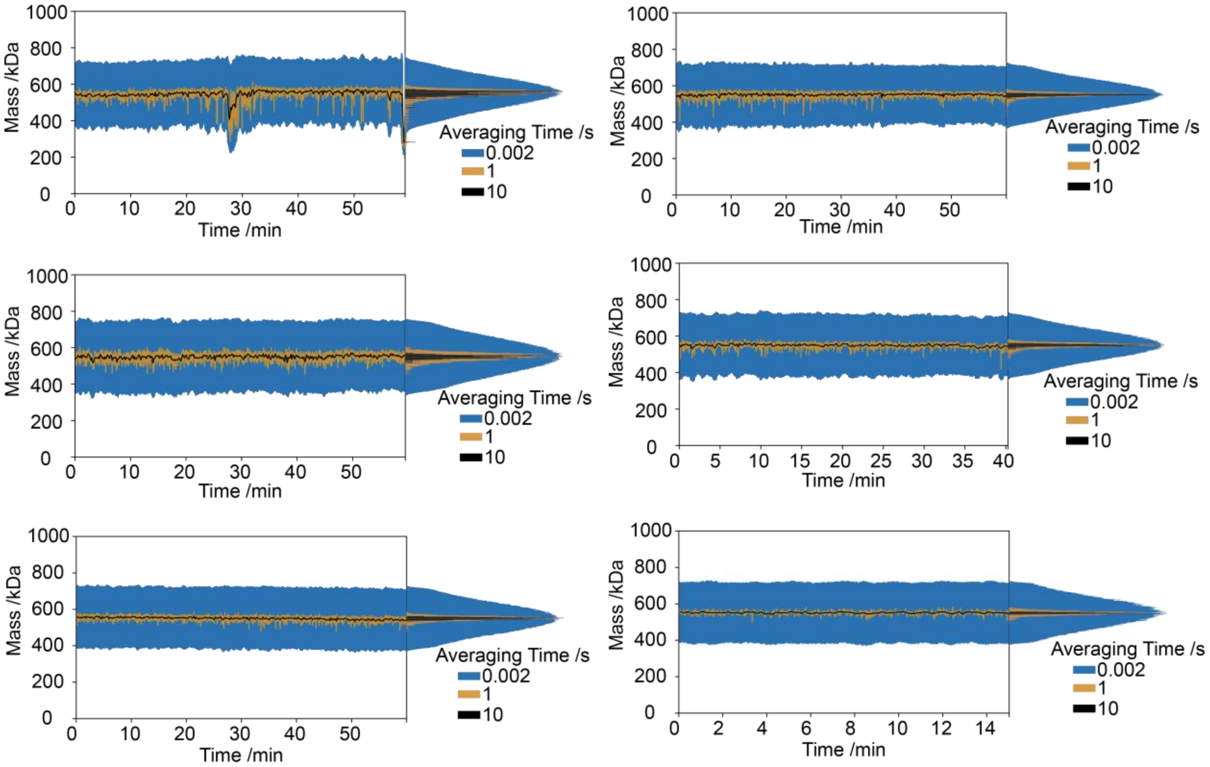

**Supplementary figure 3. SARS-CoV-2 long tracking replicates.**
Mass vs time traces of six replicates of a single SARS-CoV-2 glycoprotein diffusing across a patterned bilayer and measured with mass photometry. The patterns were of ~4 µm diameter supported lipid bilayer circles and the measurement was done with a 638 nm illumination at 497 Hz. Different averaging times show enhanced mass precision.

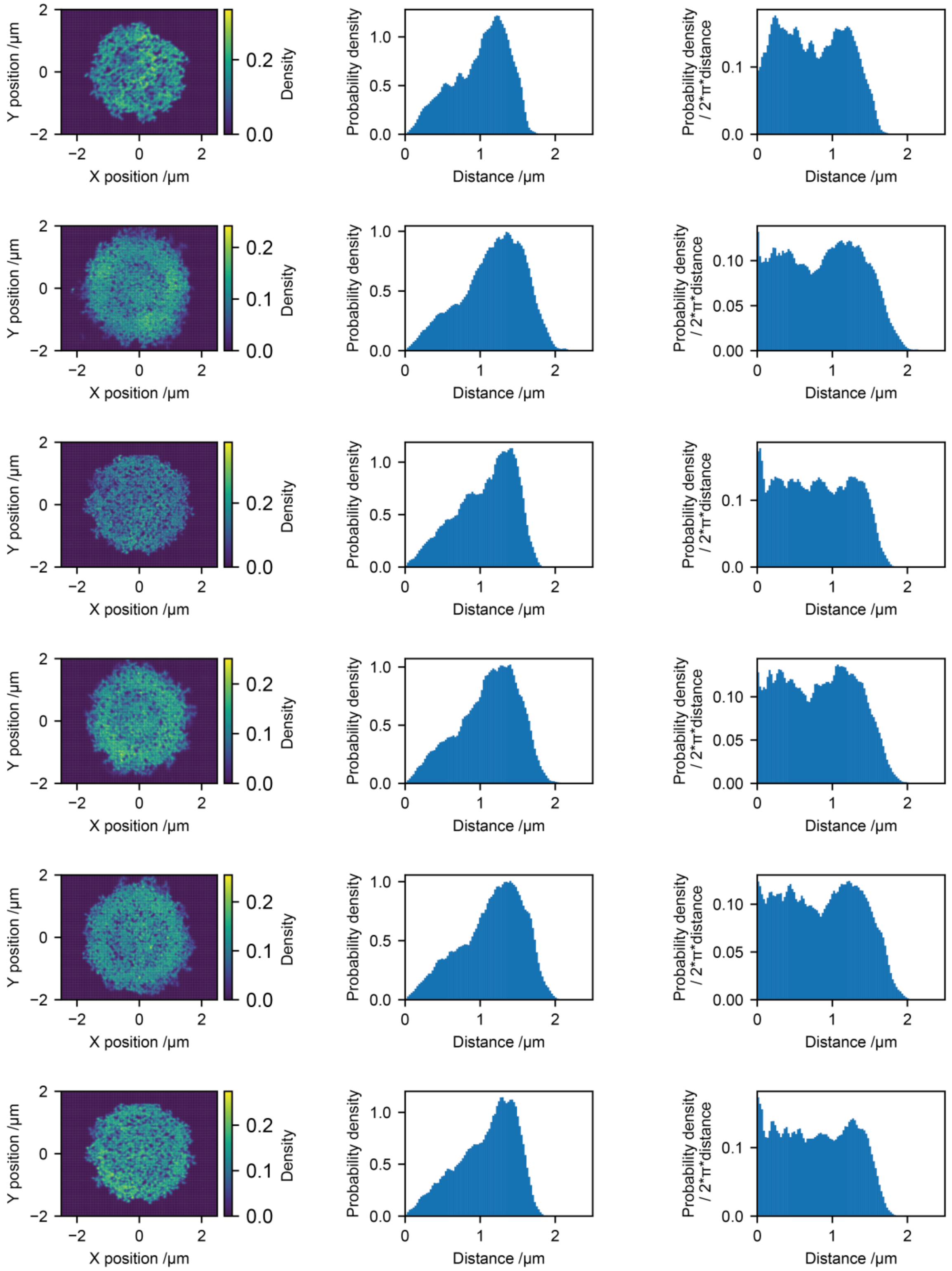

**Supplementary figure 4. Spatial distribution of confined diffusing proteins.** Left to right: Heatmap of detected SARS-CoV-2 spike locations during long tracking showing confinement to the patterned supported lipid bilayer, probability density of the detected events as a function of distance from the centre of the patterned area, normalised probability density of detected events as a function of distance from the centre of the patterned area.

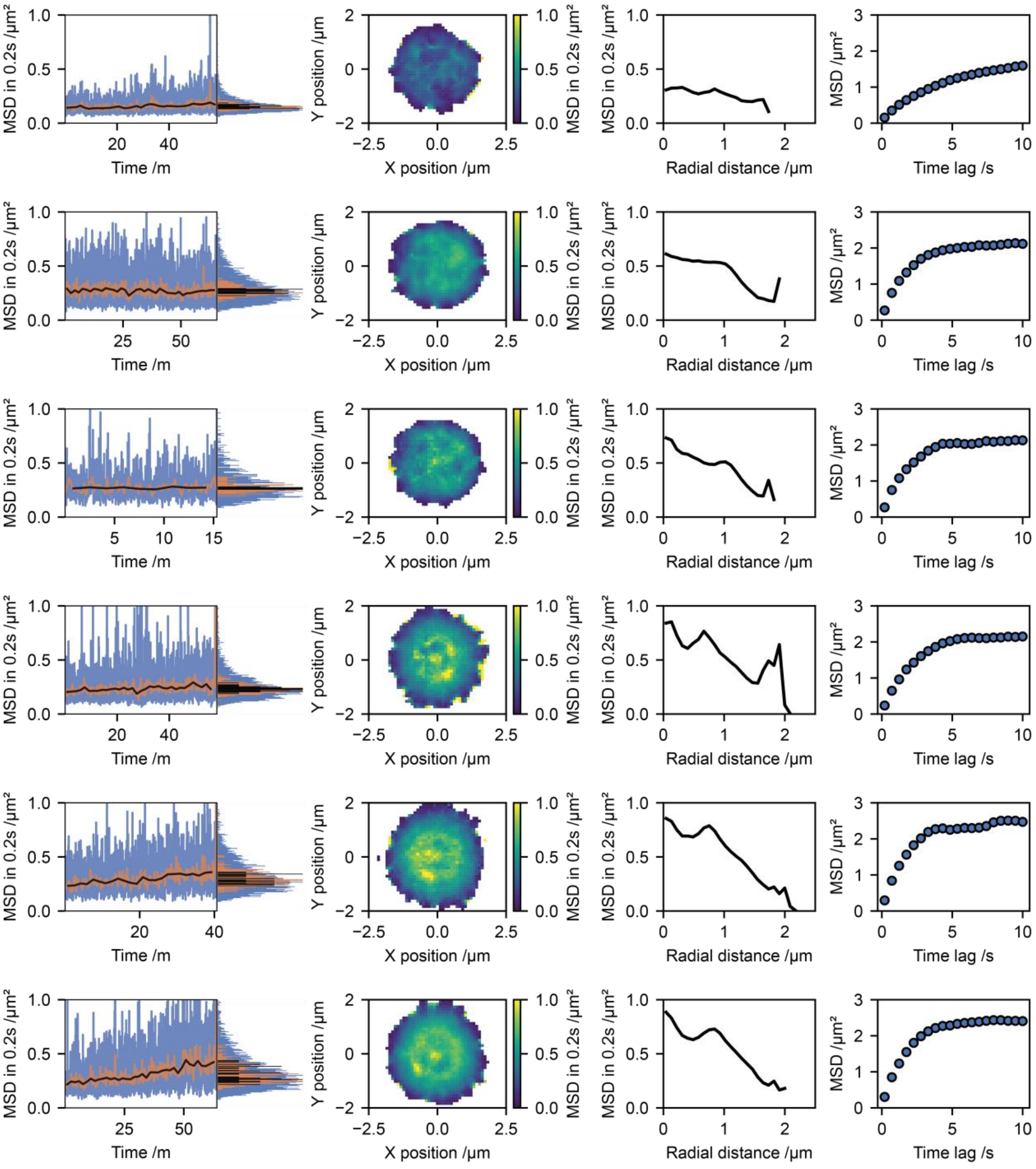

**Supplementary figure 5. Characterisation of the confined diffusion mean squared displacement (MSD).**
Left to right: Mean squared displacement (MSD) of a trapped particle in 0.2 seconds as either raw values (blue) or further time averaged for 1 second (orange) or 10 seconds (black); Heatmap of the distribution of MSD across the trap; Radial mean of observed MSD values; MSD values for different time lags.

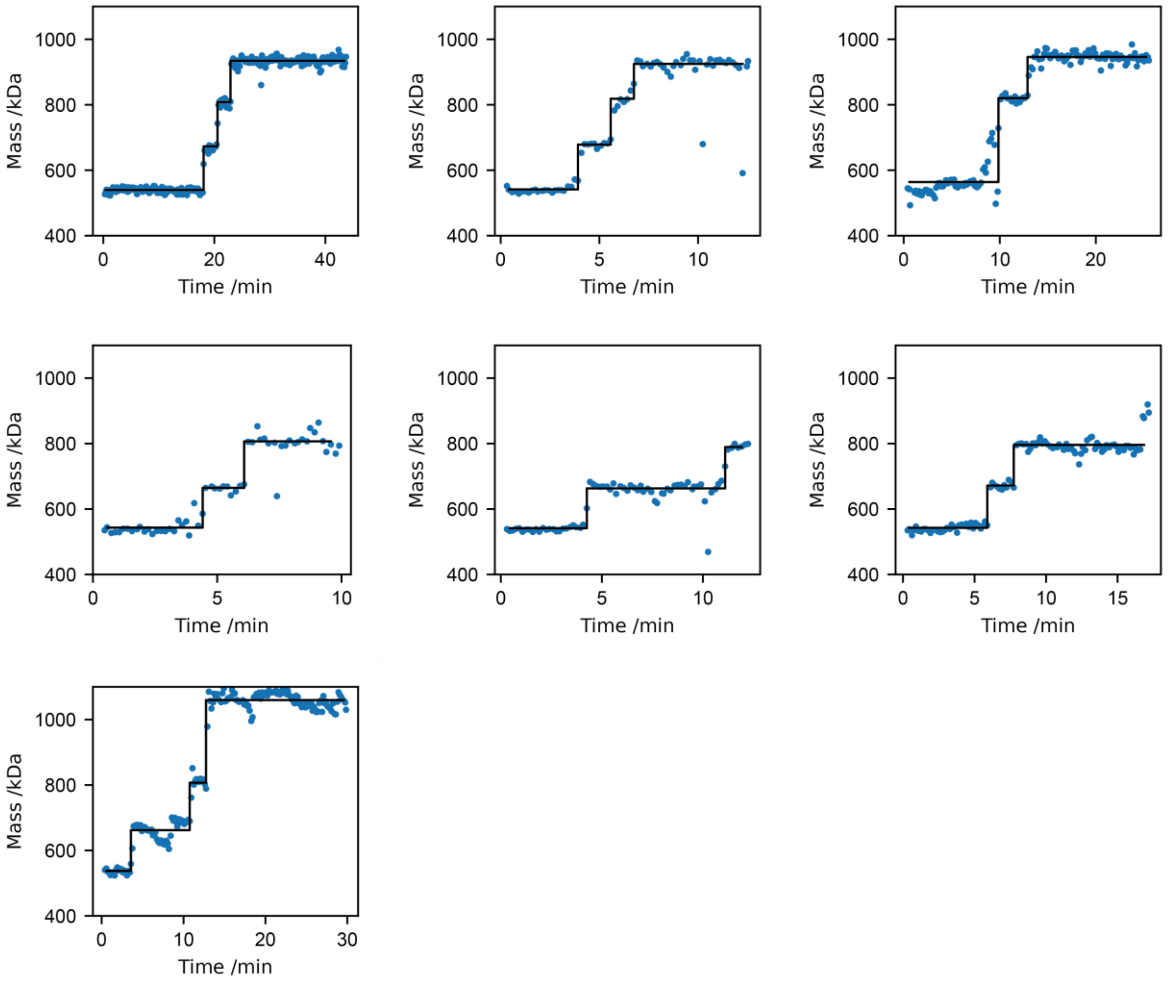

**Supplementary figure 6. Antibody binding.**
Mass vs time traces of eight replicates of stepwise antibody (150 kDa) binding to a SARS-CoV-2 spike trimer and their corresponding step function fit. Scatter points represent consecutive mass averaging windows of 10 seconds.

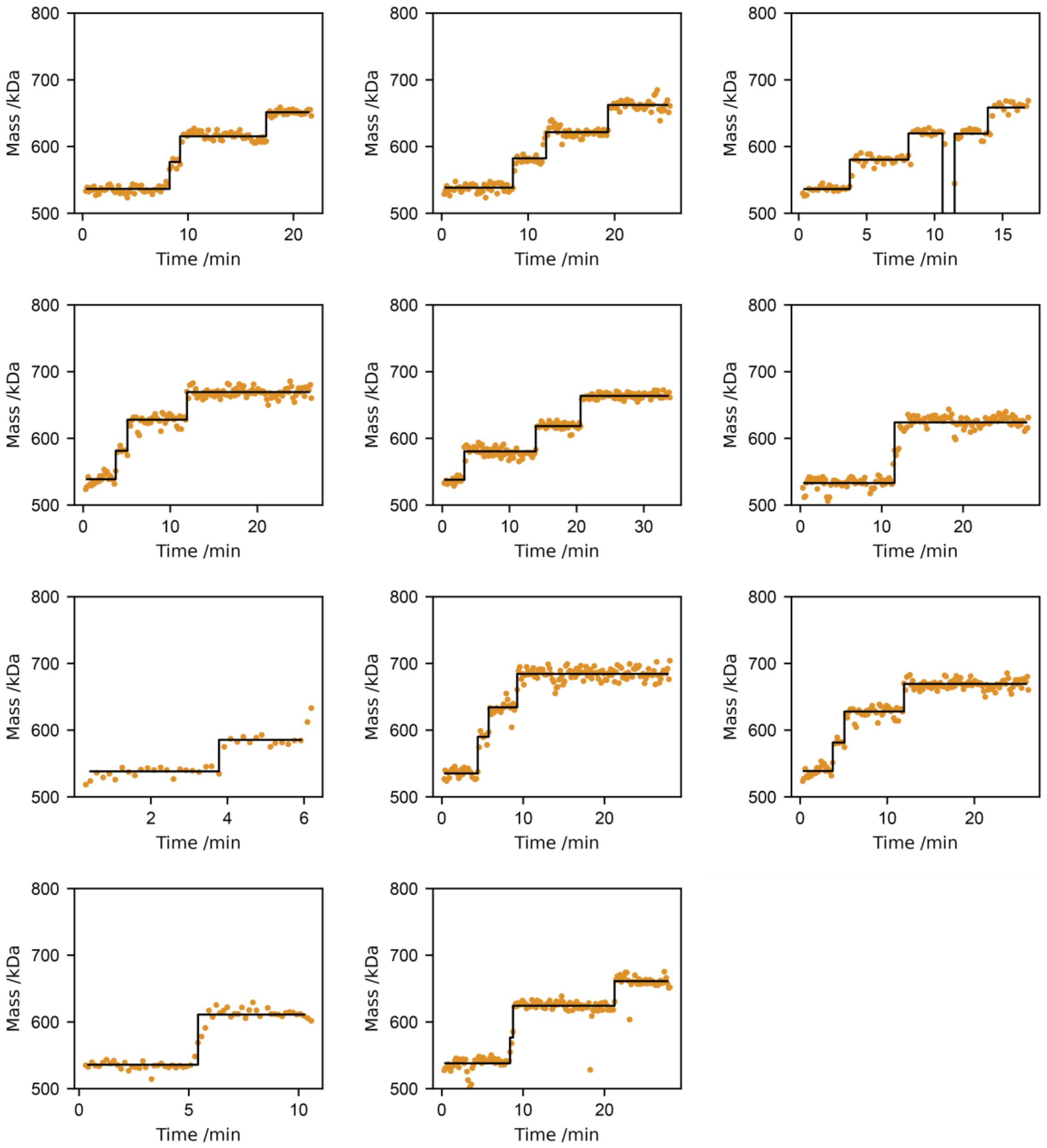

**Supplementary figure 7. Fab binding.**
Mass vs time traces of eleven replicates of stepwise Fab (49 kDa) binding to a SARS-CoV-2 spike trimer and their corresponding step function fit. Scatter points represent consecutive mass averaging windows of 10 seconds.

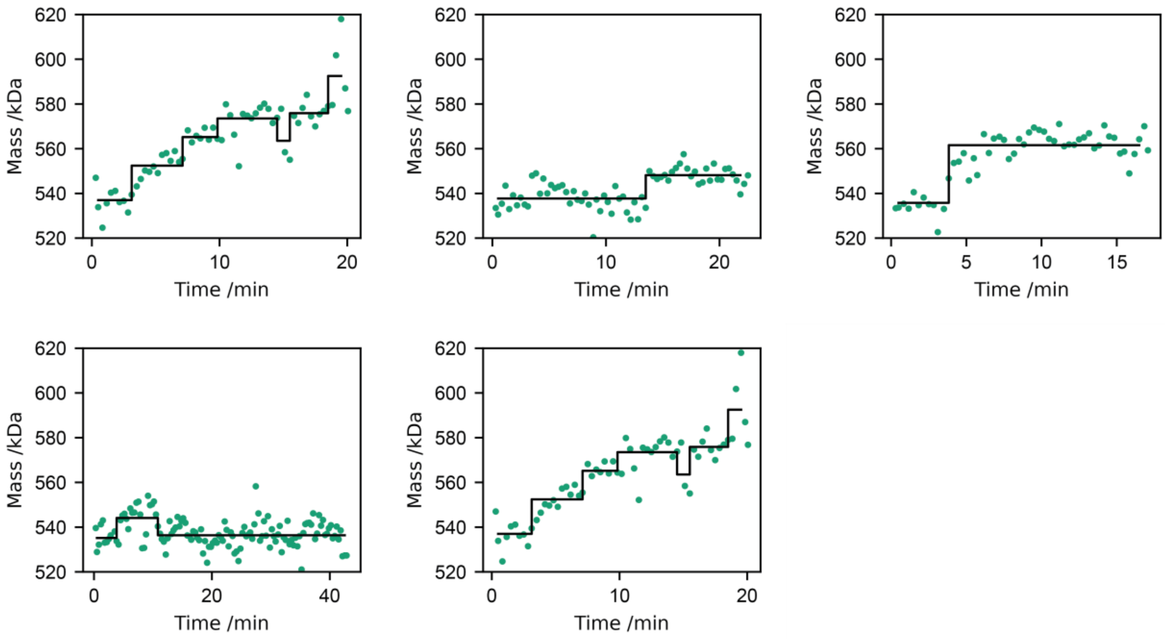

**Supplementary figure 8. Nanobody binding.**
Mass vs time traces of five replicates of stepwise nanobody (16 kDa) binding to a SARS-CoV-2 spike trimer and their corresponding step function fit. Scatter points represent consecutive mass averaging windows of 20 seconds.

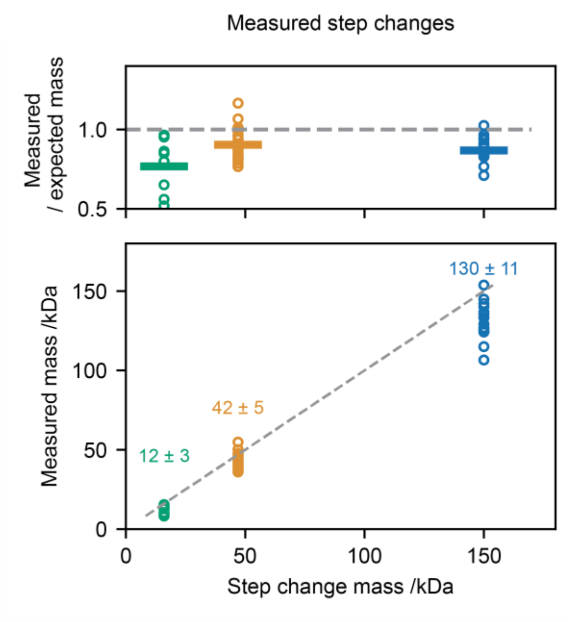

**Supplementary figure 9. Step change quantitation.**
Quantitation of the measured step changes for nanobody (green), Fab (orange) and antibody (blue) binding.

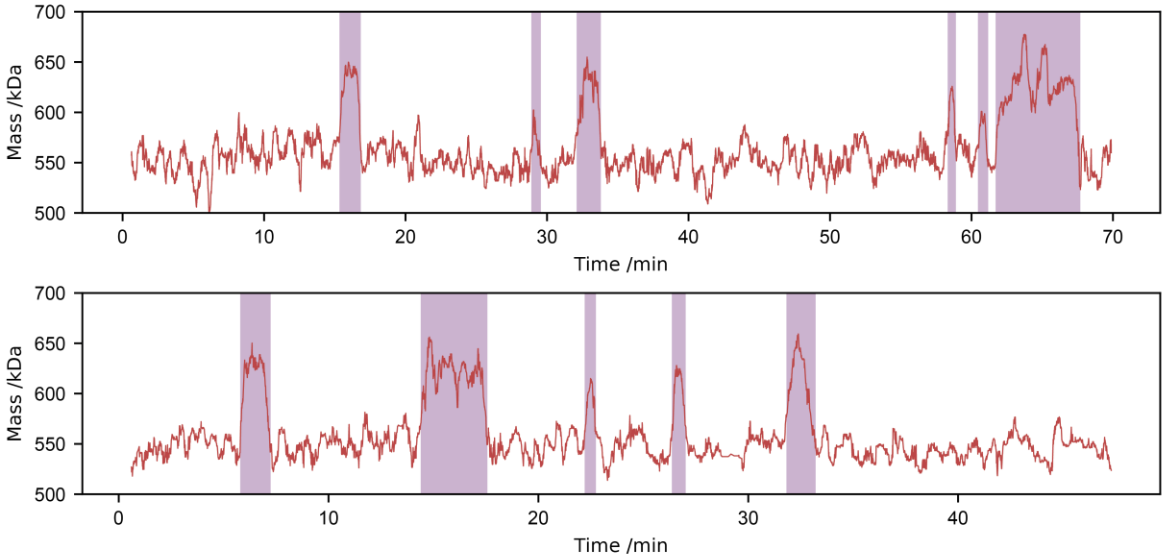

**Supplementary figure 10. Transient mACE2 binding.**
Mass vs time traces of two replicates of transient monomeric ACE2 binding to a SARS-CoV-2 spike trimer and a corresponding two state hidden Markov model fit. The red plot represents a mass averaged running median of 10 seconds and the purple areas represent the mACE2 bound state. The top plot is the same plot shown in figure 2 panels h-i.

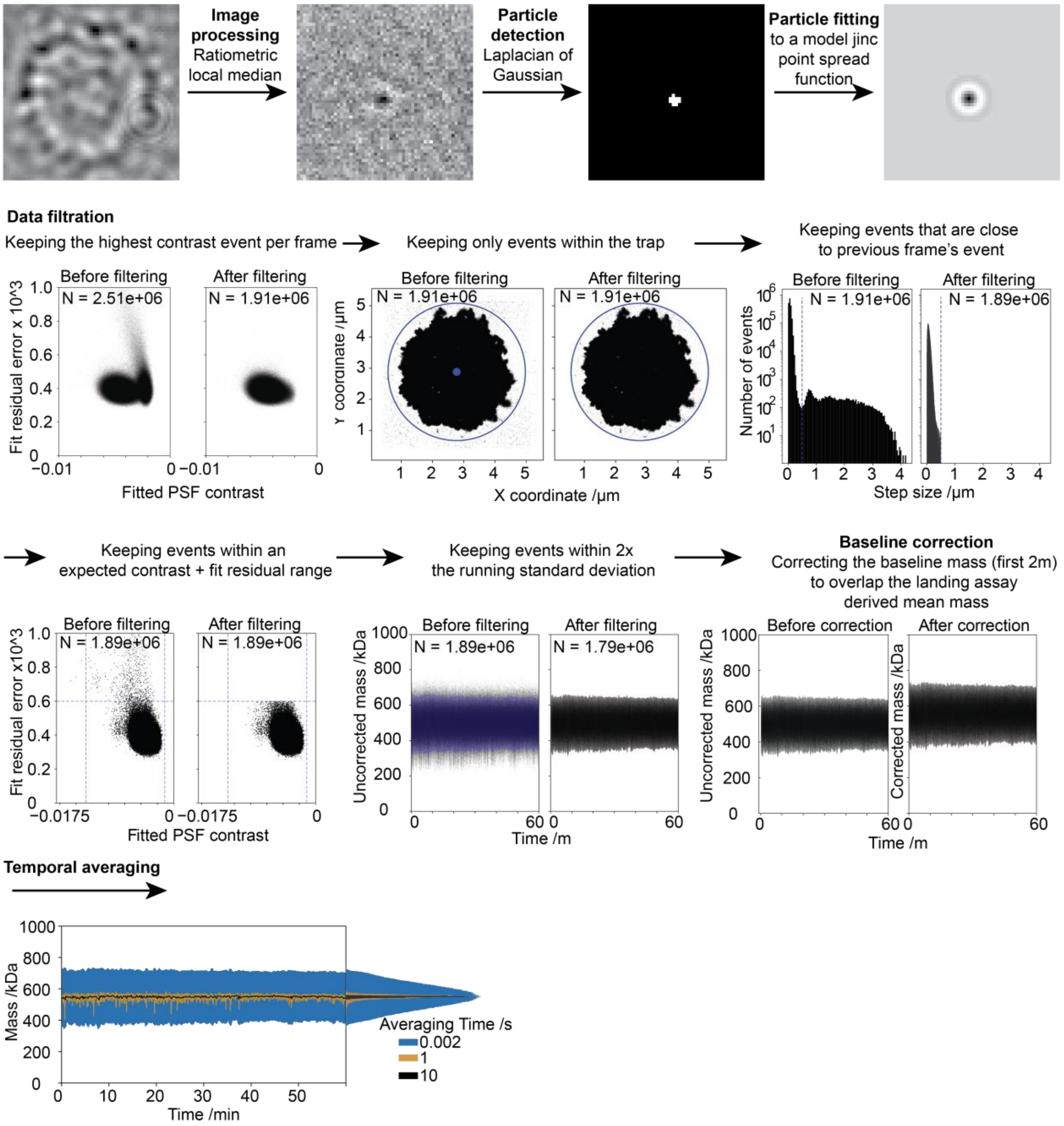

**Supplementary figure 11. Data analysis workflow.**

Schematic showing the data analysis workflow for confined diffusion mass photometry experiments starting from a raw mass photometry video up to a mass vs time trace.

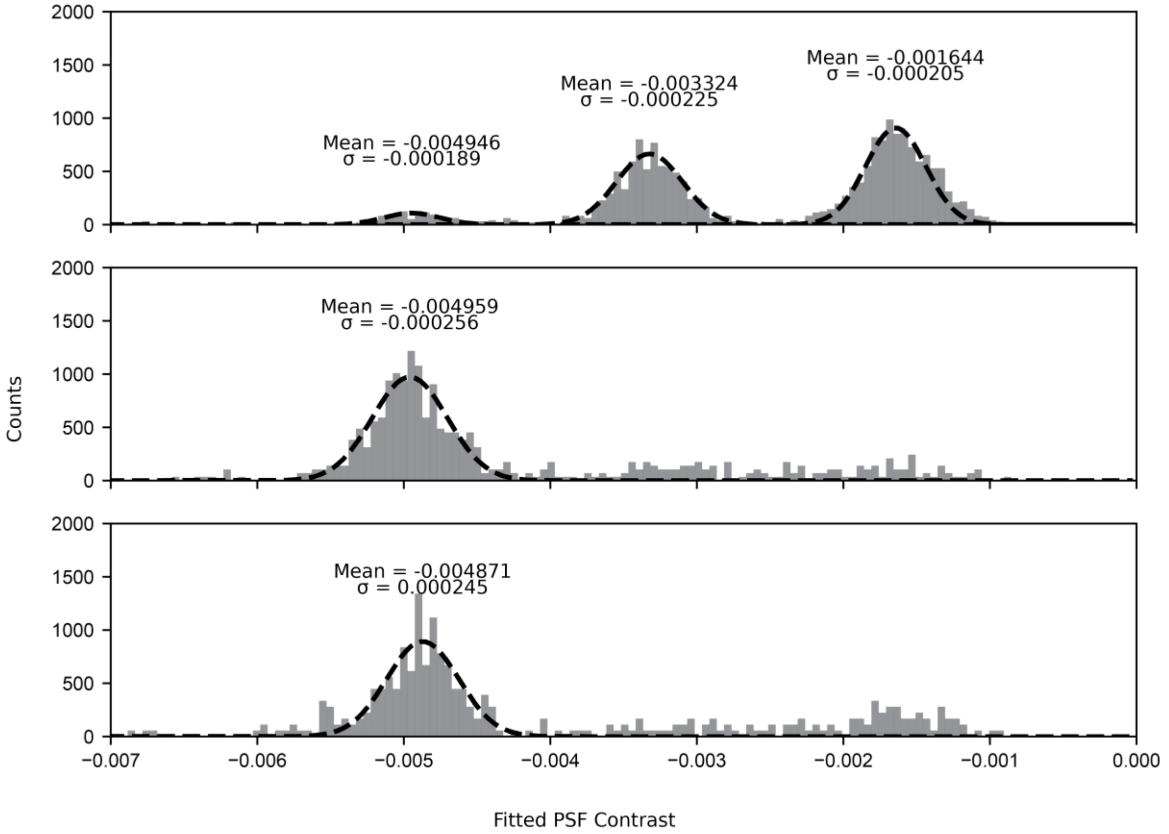

**Supplementary figure 12. Landing assay contrast to mass quantitation.**
Regular mass photometry landing assays measured at 250Hz, 2 ms exposure per frame at 638 nm and frame averaging time of 160ms for a mass calibrant (dynamin ΔPRD, top row) and SARS-CoV-2 spike (middle and bottom rows). Establishing a contrast to mass slope of -0.917E-6 [contrast/kDa] and an average SARS-CoV-2 spike mean mass of 536 kDa.

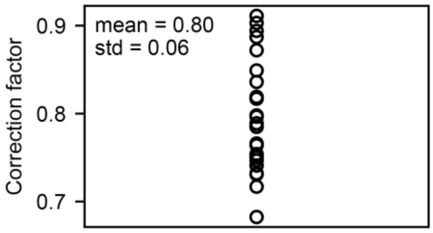

**Supplementary figure 13. Mass correction factors distribution**.
Scatter plot for the correction factors used across all of the spike traces in the manuscript in order to convert the estimated mass (fitted event contrast divided by landing assay derived mass slope) to the landing assay measured baseline mass. A correction factor of ~0.84 is to be expected, as shown previously (Asor, PNAS 2024). The replicate-to-replicate heterogeneity might stem from day-to-day mass slope differences, mass slope differences from measurement on different areas of the camera, diffusion differences in different traps and mass precision limitations given finite temporal averaging.
